# Cross-species alignment along the chronological axis reveals evolutionary effect on structural development of human brain

**DOI:** 10.1101/2024.02.27.582251

**Authors:** Yue Li, Qinyao Sun, Shunli Zhu, Congying Chu, Jiaojian Wang

## Abstract

Disentangling evolution mysteries of human brain has always been an imperative endeavor in neuroscience. On the one hand, by spatially aligning the brains between human and nonhuman primates (NHPs), previous efforts in comparative studies revealed both correspondence and difference in brain anatomy, e.g., the morphological and the connectomic patterns. On the other hand, brain anatomical development along the temporal axis is evident for both human and NHPs in early life. However, it remains largely unknown whether we can conjugate the brain development phases between human and NHPs, and, especially, what the role played by the brain anatomy in the conjugation will be. Here, we proposed to embed the brain anatomy of human and macaque in the chronological axis for enabling the cross-species comparison on brain development. Specifically, we separately established the prediction models by using the brain anatomical features in gray matter and white matter tracts to predict the chronological age in the human and macaque samples with brain development. We observed that applying the trained models within-species could well predict the chronological age. Interestingly, by conducting the cross-species application of the trained models, e.g., applying the model trained in humans to the data of macaques, we found a significant cross-species imbalance regarding to the model performance, in which the model trained in macaque showed a higher accuracy in predicting the chronological age of human than the model trained in human in predicting the chronological age of macaque. The cross application of the trained model introduced the brain cross-species age gap (BCAP) as an individual index to quantify the cross-species discrepancy along the temporal axis of brain development for each participant. We further showed that BCAP was associated with the behavioral performance in both visual sensitivity test and picture vocabulary test in the human samples. Taken together, our study situated the cross-species brain development along the chronological axis, which highlighted the disproportionately anatomical development in the human brain to extend our understanding of the potential evolutionary effects.

---

Understanding the evolutionary effects on the neural system is a fundamental objective in comparative neuroscience. Compared with nonhuman primates (NHPs), the human brain has evolved dramatically to support the human unique specialized abilities such as language, thinking, sociality, and other high-order cognitive functions (Rilling, 2014b; Saxe, 2006; Wang et al., 2020b). Due to higher similarities in brain homology, analogous socio-emotional behaviors, and genetics, the NHP, particular rhesus macaques, provide an excellent surrogate for investigating the brain evolution of the human lineage and its relationship to pathophysiology (Gray and Barnes, 2019; Howell et al., 2019; Phillips et al., 2014). Unraveling evolutionary adaptations of the human brain is not only crucial to understand the neural substrates of human unique cognitive functions but could contribute to uncovering pathological mechanisms of human brain disorders and developing effective therapeutics strategies (Feng et al., 2020; Nelson and Winslow, 2009; Thiebaut de Schotten et al., 2019).

Previous comparative studies have found that primate evolution has remodeled the brain architectures in structures, functions, and wiring patterns. With the development of macroscopic magnetic resonance imaging (MRI) technique, comparative neuroimaging could directly evaluate brain structures, functions and connectivity differences between humans and NHPs. The structural MRI revealed increased cortex expansion and hemispherical asymmetry in humans compared to NHPs (Alexander et al., 2001; Donahue et al., 2018; Hill et al., 2010; Leroy et al., 2015; Marie et al., 2018). In addition, human-specific functional brain areas or networks responsible for decision making and attention emerging during human evolution were reported (Mantini et al., 2013; Neubert et al., 2014; Patel et al., 2015). Advances in diffusion tensor imaging allow direct quantification of microstructure of fiber tracts, anatomical connectivity strength between brain areas and complex network organization between humans and NHPs. The increased myelination and fiber connections of arcuate fasciculus for language processing have been consistently found in humans compared to NHPs (Balezeau et al., 2020b; Eichert et al., 2019; Rilling et al., 2008; Sierpowska et al., 2022). Complex network analysis based on macroscale structural connectivity network revealed different network hubs, common and unique wiring properties of the human brain (Goulas et al., 2014; Li et al., 2013). Recently, we mapped evolutionary and developmental connectivity atlas of language areas and revealed increased functional balance, amplitude of low frequency fluctuations, functional integration, functional couplings and better myelination of dorsal and ventral white matter language pathways in humans compared to macaques (Cheng et al., 2021; Wang et al., 2020a). While these studies offer ample evidence of brain differences between humans and NHPs, it is important to acknowledge that quantifying evolutionary differences through spatial alignment comparisons may partially manifest the evolutionary effects on the human brain, considering the intrinsic evolutionary divergence between humans and macaques about 25 million years ago.

Except for the spatial alignment between the cross-species brains, mounting evidence has demonstrated the existence of brain development along the temporal axis for both humans and macaques in the early life. Studies on early human brain development have shown that the sensory cortex tends to develop more quickly than the association cortex, and that language processing specialization occurs in the left hemisphere even during infancy (Dubois et al., 2014; Paus et al., 2001). Furthermore, cognitive control, emotional processing, and motivation are interdependent and mature during adolescence (Christakou, 2014; Cunningham et al., 2002; Kilford et al., 2016). Additionally, the cortical surface area expands during puberty in rhesus macaques (Ronan et al., 2014). Upon the phenomenon of the co-existence of brain development, it raises the question whether we can conduct the cross-species comparison on the brain by comparing the phases of brain development. In other words, it would be necessary to establish a comparable relationship between brain anatomy and the phases of brain development. By doing so, it would add our understanding: 1). Differences in the conserved and dominant of human and macaque brain structure, function, and networks at corresponding developmental stages. 2). Directly quantify the disproportionate anatomical development and evolution in human and macaque brain at specific developmental stages.

To address the current research bottlenecks of comparative neuroscience lacking analyzing human and macaque developmental data along a chronological axis. We here proposed a machine learning based cross-species prediction model, i.e., brain structure based cross-species age prediction model, to quantitatively characterize brain evolutionary pattern along the temporal axis. First, the gray matter volume (GMV) and microstructure of white matter tracts (fractional anisotropy (FA), mean diffusion (MD), axial diffusion (AD), and radial diffusion (RD)) were taken as features to train the prediction models and were used to predict ages for human and macaque, respectively. Then, the trained human and macaque prediction models were separately applied to predict the ages of macaque and human. Next, based on the macaque prediction model to predict human ages, we proposed a new concept of the brain cross-species age gap (BCAP) to quantify evolutionary differences. Finally, Pearson’s correlations were performed to identify BCAP associated brain areas, white matter tracts and behavioral phenotypes. The details of the pipeline of this study are shown in Figure 1.

**Figure 1.**
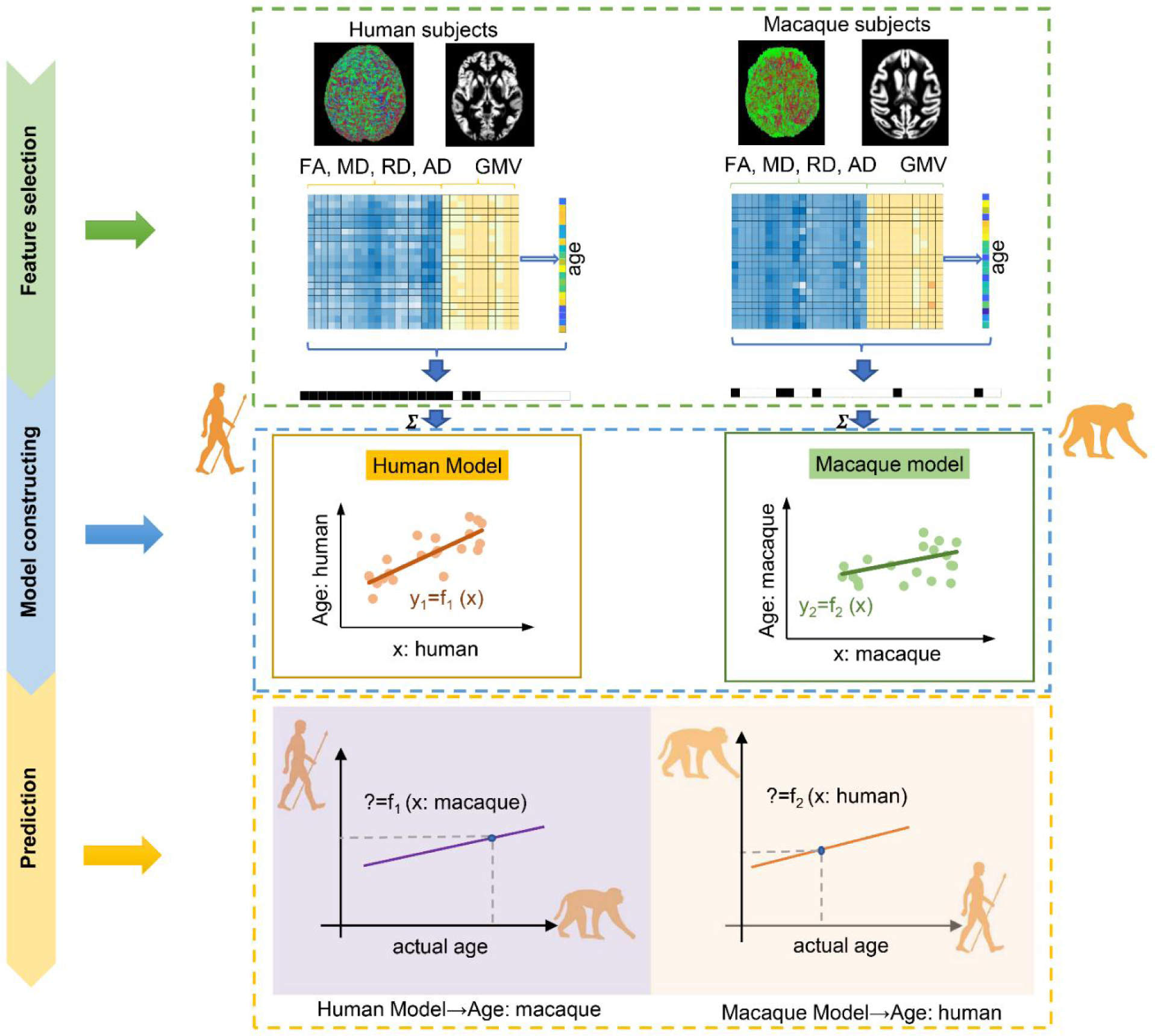
The flowchart for intra- and inter-/cross-species prediction using brain structure based cross-species age prediction model. Feature selection: the features with *p* < 0.01 from Pearson’s correlation between features consisted of gray matter volume (GMV), fractional anisotropy (FA), mean diffusion (MD), radial diffusion (RD) and axial diffusion (AD) and ages were preliminarily selected. The ultimate features were selected using two criterions: common features and minimum mean absolute error (MAE) across 100 times repetitions. Model construction: using the selected features, 10-fold cross-validation with nine tenths was used to train model. Prediction: 10-fold cross-validation with one tenth was used for prediction.

## Results

### Intra- and inter-/cross-species age prediction

The humans and macaques brain ages were predicted using brain structure based cross-species age prediction model. First, we used 62 macaque features and 225 human features which were present in all the 100 times iterations to train the macaque and human prediction models, respectively (all the features see Table supplement 1 and 2 in Supplementary information). Using the trained models, we found that the macaque prediction model can well predict macaque ages (R = 0.5729, P < 0.001, MAE = 0.3758; Figure 2(A)) and the human prediction model can well predict human ages (R = 0.6153, P < 0.001, MAE = 1.1236; Figure 2(D)). Interestingly, we applied the trained macaque and human prediction models to separately predict human and macaque ages for inter-/cross-species prediction. We found that the trained monkey prediction model could well predict human ages (R = 0.4823, P < 0.001, MAE = 8.3610; Figure 2(B)), and the trained human prediction model can also predict macaque ages (R = 0.2898, P < 0.001, MAE = 7.6157; Figure 2(C)). However, we noticed that using the macaque prediction model to predict human ages showed better performance than using the trained human prediction model to predict macaque ages.

**Figure 2.**
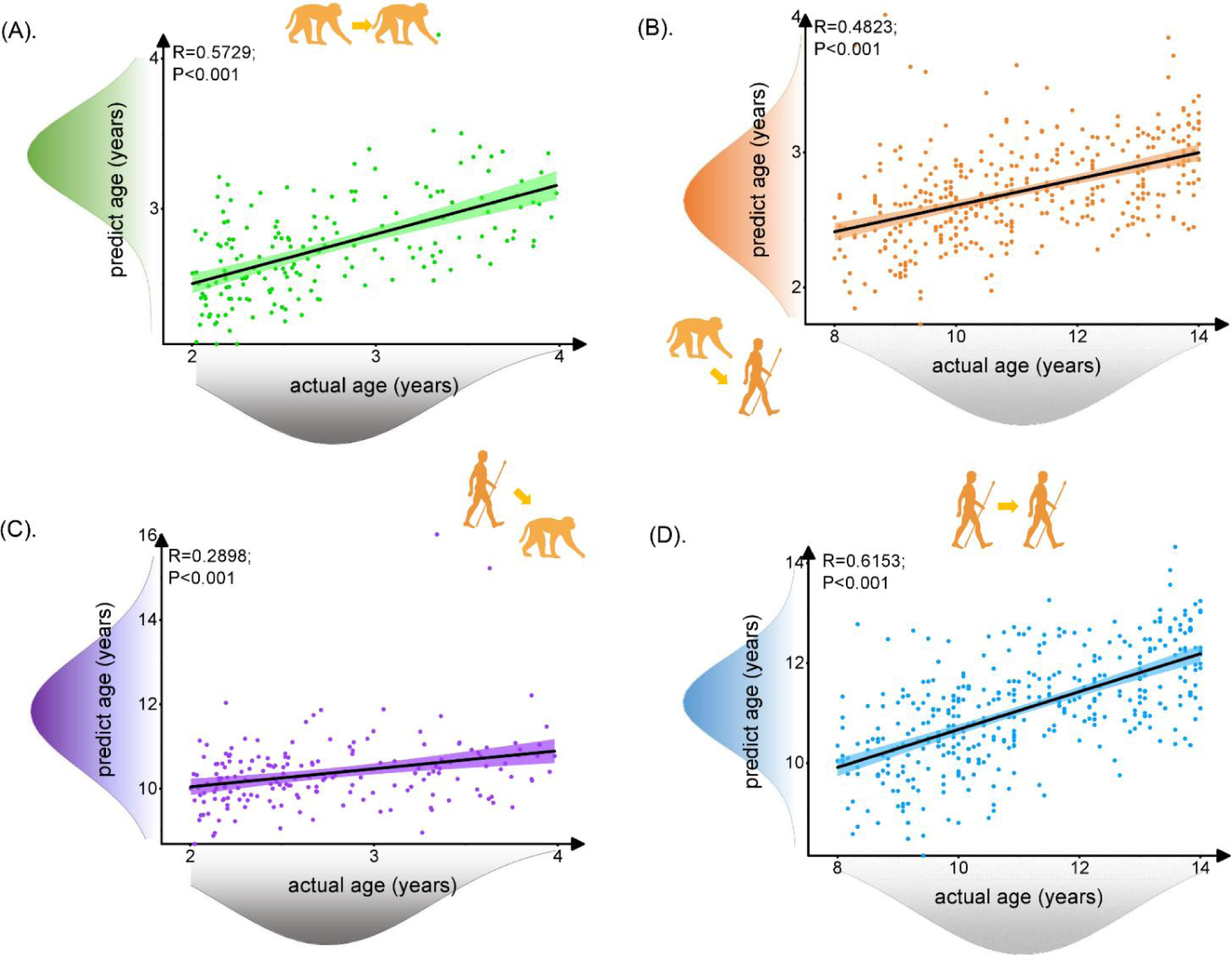
The prediction results for intra- and inter-/cross-species using brain structure based cross-species age prediction model. Each dot depicts data from an individual participant. The width of the curve denotes the 95% confidence interval around the linear fitting curve (black line). The prediction model could well predict intra- and inter-species ages. The trained monkey prediction model could better predict human ages than using trained human prediction model to predict macaque ages. (A), (D). Prediction results for intra-species with green for macaque and blue for human. (B), (D). Prediction results for inter-species with orange for predicted human ages using macaque model and purple for predicted macaque ages using human model.

To test the impact of different number of features on prediction performance, we first trained the macaque prediction model with 117 features and human prediction model with 239 features selected using the criterion of minimum MAE during model training with 100 repetitions (all the features see Table supplement 3 and 4 in Supplementary information). We found that the macaque prediction model and human prediction model can well predict macaque (R = 0.5825, P < 0.001, MAE = 0.3675) and human ages (R = 0.6039, P < 0.001, MAE = 1.1388), respectively. We also found that the trained monkey prediction model could well predict human ages (R = 0.4018, P < 0.001, MAE = 7.7185) and the trained human prediction model can also predict macaque ages (R = 0.3223, P < 0.001, MAE = 7.9514) (Figure supplement 1 in Supplementary information).

Using the same top 62 features of macaque and humans, we observed that the macaque prediction model and human prediction model can well predict macaque (R = 0.5729, P < 0.001, MAE = 0.3760) and human ages (R = 0.6818, P < 0.001, MAE = 1.0025), respectively. We also observed that the trained macaque prediction model could well predict human ages (R = 0.4822, P < 0.001, MAE = 8.3606) and the trained human prediction model can also predict macaque ages (R = 0.2094, P = 0.0047, MAE = 6.1083). Through testing different number of features, we found that both human and macaque prediction models could well predict their corresponding ages. In addition, we also observed a good performance for the inter/cross-species prediction. Consistently, we showed that the trained macaque prediction model predicting human ages outperformed the trained human prediction model predicting macaque ages even using different number of features for inter-/cross-species prediction (Figure supplement 2 in Supplementary information).

Finally, to explore whether sex affects the prediction model, we separated both human and macaque participants into males and females’ groups for intra- and inter-species prediction. The predicted results showed similar pattern with that predicted by combining male and female into one group (Figure supplement 3 in Supplementary information).

By testing different number of features and sex effects, we observed consistent patterns for intra- and inter-/cross-species age prediction using prediction model. Importantly, we revealed a stable pattern that the trained macaque prediction model could better predict human ages than that using the trained human prediction model to predict macaque ages.

### Ages effects on the brain age gap of |Δ _brain age_| for the inter-/cross-species prediction

We evaluated the relationship between actual age and predicted age deviation (|Δ _brain age_|) for inter-/cross-species prediction. When we used macaque model to predict human ages, the |Δ _brain age_| in human showed a positive correlation with actual age of the human subjects indicating that the error of using macaque model to predict human age increased while human age increases. On the contrary, when using human model to predict macaque age, the prediction error of the human model decreased as the macaque age increases (Figure 3).

**Figure 3.**
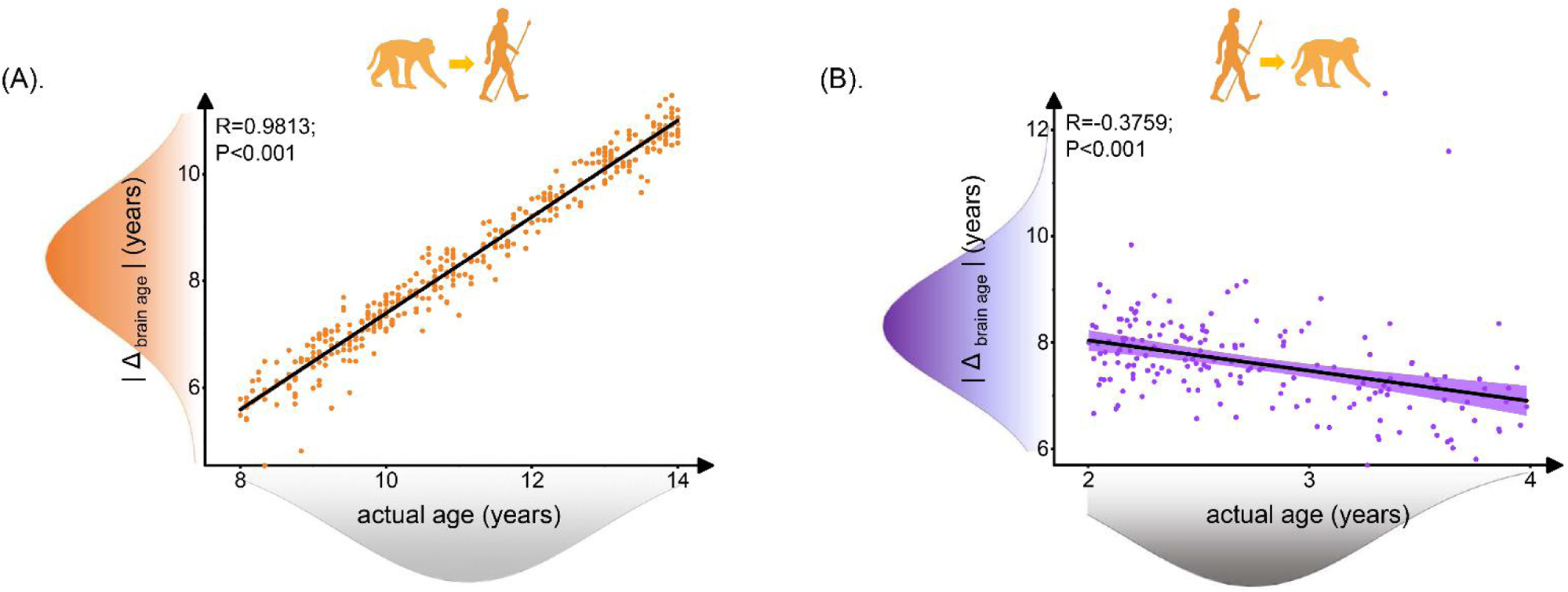
Relationship between brain age gap (|Δ _brain age_|) and actual ages in human. Each dot depicts data from an individual participant. The width of the curve denotes the 95% confidence interval. The human brain age gap (|Δ _brain age_|) is defined as predicted human age using macaque model minus human actual age. The macaque brain age gap (|Δ _brain age_|) is defined as predicted macaque age using human model minus macaque actual age. (A). Positive association between |Δ _brain age_| and actual age in human ages was found (Pearson’s correlation: R = 0.9813, P < 0.001, MAE = 2.7120). (B). Negative association between | Δ _brain age_ | and actual ages in macaque was found (Pearson’s correlation: R = −0.3759, P < 0.001, MAE = 4.8697).

Like above, we also conducted the above correlation analyses between actual ages and predicted age deviation using 117 macaque features and 239 human features and the same top 62 features in both macaque and human, the results obtained with the two sets of features showed the similar patterns demonstrating the stability of the prediction models (Figure supplement 4 and 5 in Supplementary information). In addition, to test sex effects, the trends of the correlations between actual age and predicted age deviation are similar with that found using different sets of features suggesting the results had no significant difference between sexes (Figure supplement 3 in Supplementary information). The decreased prediction accuracy using macaque prediction mode for human ages suggests growing evolutionary differences during human development.

### The brain areas and white matter tracts contributing to prediction

We classified the features for predication in human and macaque into three subtypes (human-specific features, macaque-specific features and common features of humans and macaques) and the proportion of each subtype of features belonged to GMV, FA, MD, AD and RD was calculated. The top five brain areas or white matter tracts showing highest correlations with ages were displayed (Figure 4). We observed that macaque-specific features were mainly located in gray matter areas, while human-specific features were mainly distributed in white matter tracts and only small proportions in gray matter areas. The common features of human and macaque have the highest proportion in gray matter areas but also some were located in white matter tracts (Figure 4(A)). For the macaque-specific features, the gray matter features are mainly located in the left dorsolateral prefrontal cortex (PFCdl, R = 0.4003), left medial prefrontal cortex (PMCm, R = 0.3966), right dorsolateral prefrontal cortex (PFCdl, R = 0.3723), right orbitomedial prefrontal cortex (PFCom, R = 0.3478) and right centrolateral prefrontal cortex (PFCcl, R = 0.3427) and the white matter features are mainly located in the left superior longitudinal fasciculus II (SLF II, R = 0.3067) and right uncinate fasciculus (UF, R = 0.2901) (Figure 4(B)). For the human-specific features, the gray matter features are mainly located in left Putamen (Pu, R = 0.4437), right Pallidum (GP, R = 0.4284), left posterior insula (Ip, R = 0.3915), left Pallidum (GP, R = 0.3884) and right primary somatosensory cortex (S1, R = 0.3703) and the white matter features are mainly located in the right superior longitudinal fasciculus I (SLF I, R = 0.4138), left inferior fronto-occipital fasciculus (IFOF, R = 0.3665), left arcuate fasciculus (AF, R = 0.3459), right inferior fronto-occipital fasciculus (IFOF, R = 0.3445) and left superior longitudinal fasciculus I (SLF I, R = 0.3443) (Figure 4(C)). The common features of human and macaque are also distributed in gray matter and white matter tracts. The top 5 brain areas of common gray matter features in human are right posterior insula (Ip, R = 0.3769), left posterior cingulate cortex (CCp, R = 0.3583), left ventrolateral premotor cortex (PMCvl, R = 0.3523), right posterior cingulate cortex (CCp, R = 0.3466) and right intraparietal cortex (PCip, R = 0.3433) while the top 5 brain areas of common gray matter features in macaque are right anterior cingulate cortex (CCa, R = 0.4694), left anterior cingulate cortex (CCa, R = 0.4274), right medial premotor cortex (PMCm, R = 0.4235), left superior parietal cortex (PCs, R = 0.4228) and right inferior parietal cortex (PCi, R = 0.4035). The top 5 common white matter tract features in human were left corticospinal tract (CT, R = 0.4654), right corticospinal tract (CT, R = 0.4499), right superior thalamic radiation (STR, R = 0.3344), left superior thalamic radiation (STR, R = 0.3212) and left cingulum subsection: peri-genual (Cs:Pg, R = 0.2341), and the top 5 common white matter tract features in macaque are right corticospinal tract (CT, R = 0.4189), left corticospinal tract (CT, R = 0.3746), right uncinate fasciculus (UF, R = 0.3497), left superior longitudinal fasciculus II (SLF II, R = 0.3067) and left superior thalamic radiation (STR, R = 0.3046) (Figure 4(D)).

**Figure 4.**
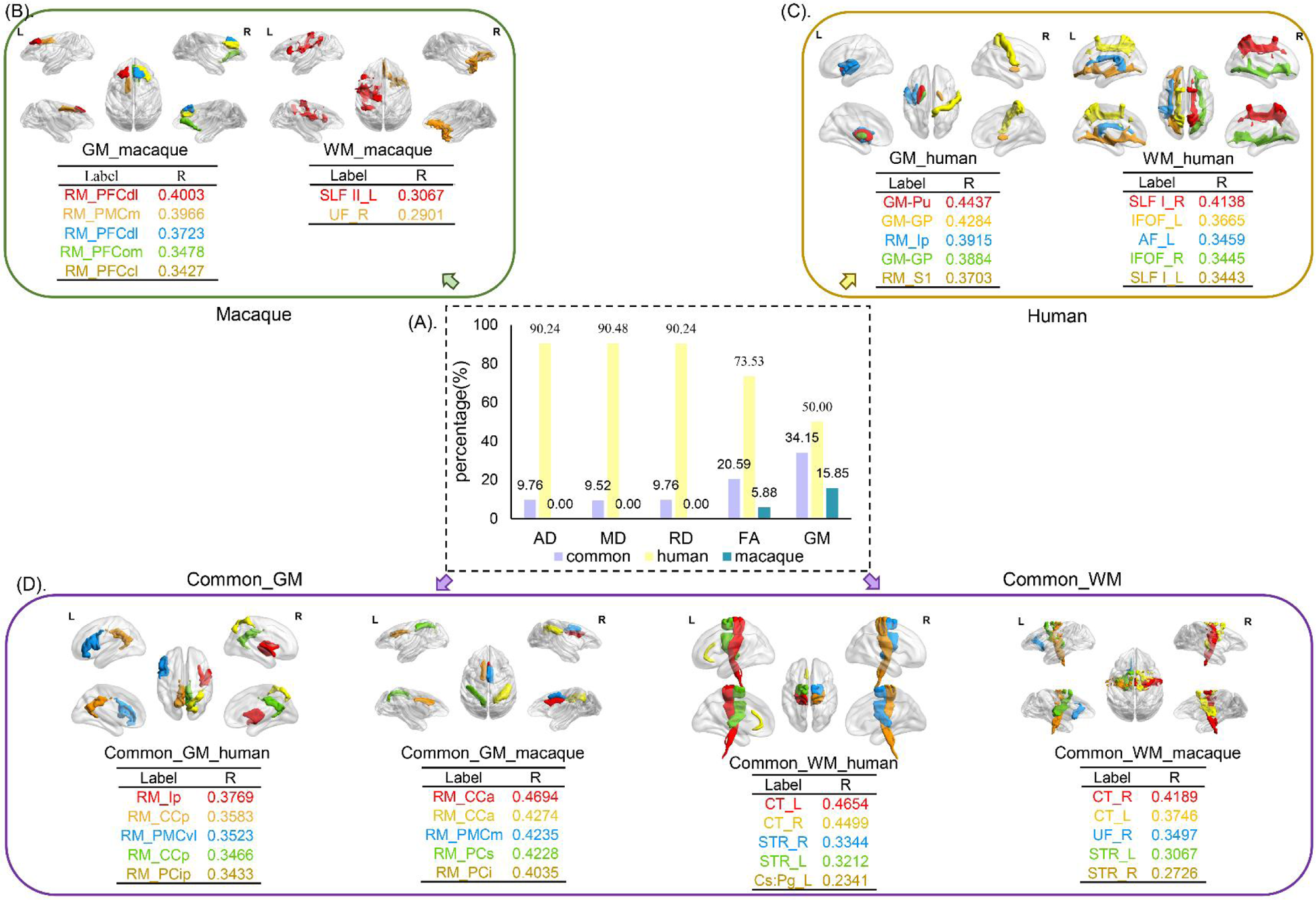
The distribution of selected features for prediction. The selected features were analyzed based on five parameters (GMV, FA, MD, RD and AD) and three groups (human-specific, macaque-specific and common features in human and macaque). (A). The percentage of each group in each parameter. The macaque-specific features are only located in FA and GMV. (B), (C), (D). The top five features of macaque-specific, human-specific, and common in human and macaque in gray matter and white matter tracts were shown. FA: fractional anisotropy; MD: mean diffusivity; RD: radial diffusivity; AD: axial diffusivity; GMV: gray matter volumes.

Meanwhile, using 117 macaque features and 239 human features or the same top 62 features of human and macaque for intra- and inter-/cross-species predication, we observed the distribution of the human-specific, macaque-specific and common features in gray matter brain areas and white matter tracts are similar with the above results obtained with 62 macaque features and 225 human features derived prediction models. However, most of the macaque-specific features are located in gray matter while only a few human-specific features are located in gray matter. The highly correlated common feature in human is corticospinal tract (CT) while the highly correlated common macaque feature is anterior cingulate cortex (CCa) (Figure supplement 6 and 7 in Supplementary information).

### The brain cross-species age gap associated behavioral phenotypes, gray matter and white matter tracts

We explored the behavioral significances of BCAP using correlation analyses between behavioral scales and BCAP. We observed that BCAP shows significant correlations with picture vocabulary test (R = 0.1588; P = 0.0323) and visual sensitivity test (R = − 2051; P = 0.0056) (Figure 5(A)).

**Figure 5.**
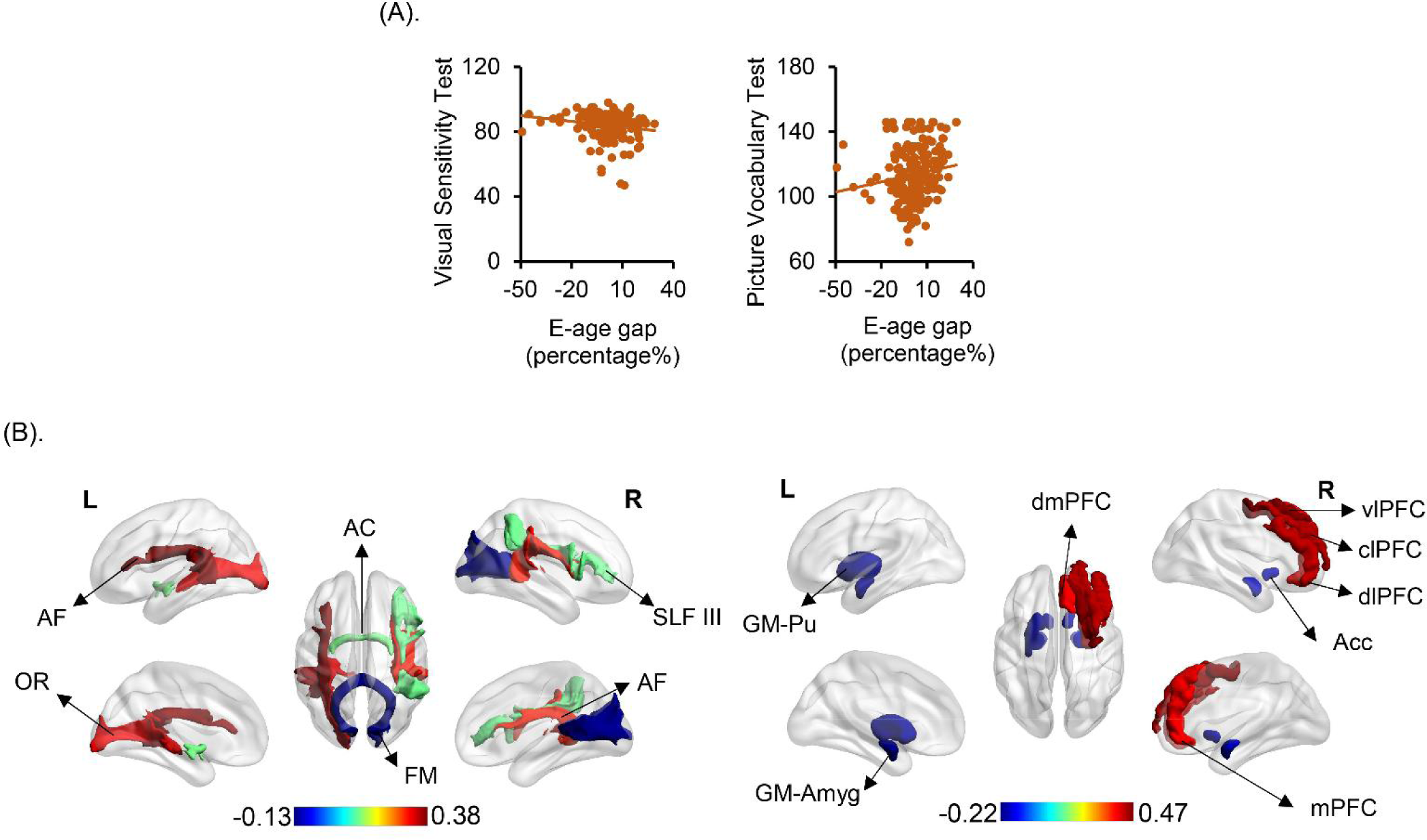
The brain cross-species age gap (BCAP) correlations with behavioral phenotypes, gray matter, and white matter tracts. (A). Each dot depicts data from an individual participant. Visual Sensitivity Test and Picture Vocabulary Test showed negative and positive correlations with BCAP, respectively. (B). The top three and last three features associated to BCAP in white matter tracts on the left, and top five and last five features associated to BCAP in gray matter on the right. These white matter tracts and gray matter related to the language pathways, emotion, and higher-order cognitive functions. AF: arcuate fasciculus; OR: optic radiation; AC: anterior commissure; FM: forceps major; SLF III: superior longitudinal fasciculus III; Pu: Putamen; Acc: Accumbens nucleus; Amyg: Amygdala; dmPFC: dorsomedial prefrontal cortex; vlPFC: ventrolateral prefrontal cortex; mPFC: medial prefrontal cortex; dlPFC: dorsolateral prefrontal cortex; clPFC: centrolateral prefrontal cortex.

Finally, with our proposed BCAP index, we identified evolution associated brain areas and white matter tracts in human. The first three and last three white matter tracts in FA showing significantly positive and negative correlations with BCAP were: left arcuate fasciculus (AF, R = 0.3784), left optic radiation (OR, R = 0.3232), right arcuate fasciculus (AF, R = 0.3035) and anterior commissure (AC, R = 0.1215), right superior longitudinal fasciculus III (SLF III, R = 0.1166), forceps major (FM, R = −0.1221); The first five and last five gray matter brain areas showing significantly positive and negative correlations with BCAP were: right dorsolateral prefrontal cortex (PFCdl, R = 0.4685), right centrolateral prefrontal cortex (PFCcl, R = 0.4324), right ventrolateral prefrontal cortex (PFCvl, R = 0.4287), right dorsomedial prefrontal cortex (PFCdm, R = 0.3915), right medial prefrontal cortex (PFCm, R = 0.3890) and right amygdala (Amyg, R = −0.1759), right accumbens nucleus (Cd, R = −0.181), left accumbens nucleus (Cd, R = −0.1811), left putamen (Pu, R = −0.2104), left amygdala (Amyg, R = −0.2151) (Figure 5(B)). The first three and last three white matter tracts between BCAP and MD, RD, and AD see in Figure S8 in Supplementary information.

## Discussion

In this study, we performed cross-species comparisons of brain development by embedding human and macaque brain anatomy in the perspective of chronological axis applying brain structure based cross-species age prediction model using multimodal brain imaging data of sMRI and dMRI to quantitatively characterize brain evolutionary patterns. We observed that the prediction model had good performances for intra-/inter-cross-species age predictions regardless of the number of features and sex. Interestingly, we found the imbalanced model performance in inter-species, which was the trained macaque prediction model to predict human ages had higher accuracy than using the trained human prediction model to predict macaque ages. Based on prediction model, we proposed the concept of the brain cross-species age gap (BCAP) index to quantify the cross-species divergence along the temporal axis during the development and further revealed that BCAP was closely associated with visual sensitivity test and picture vocabulary test indicating its potential behavioral significance. Finally, we used BCAP to investigate the evolution along with chronological axis and found that it was mainly associated with language related white matter pathways and high-order cognitive functions. Taken together, we studied the cross-species brain development along with chronological axis to quantify the disproportionately anatomical development in the human brain, which may provide a new avenue for evolutionary research of the comparative neuroscience.

### Brain structure based cross-species age prediction model and the brain cross-species age gap

The traditional comparative studies investigated brain evolution using statistical analyses by spatially aligning the brains between humans and nonhuman primates (Balezeau et al., 2020b; Gabi et al., 2016; Goulas et al., 2014; Rilling, 2014a; Wang et al., 2020a). Given the intrinsic evolutionary differences of human compared to other nonhuman primates, whether the identified differences could reflect the functional evolution along time axis remains controversial. In this study, we developed a new approach of brain structure based cross-species age prediction model and a novel perspective of chronological axis to investigate brain development and evolution. Prediction model could well predict ages for intra-and inter-species regardless of the number of features and sex effect demonstrating the reliability and stability of the developed method. For the cross-species prediction, we find that the brain age gap (i.e., |Δ _brain age_|) between human actual ages and predicted human age using macaque prediction model showed positive correlations with human actual age while the macaque brain age gap (i.e., |Δ _brain age_|) between macaque actual ages and predicted macaque ages using human prediction model showed negative correlations with the macaque actual ages, which indicated that macaque prediction model could better predict the ages of young human participants while the human macaque prediction model could better predict the ages of the older macaques. By analyzing the prediction results corresponding to human and macaque along the timeline, we speculated that the early developed brain functions are more conservative than the late developed high-order cognitive functions during species brain evolution. The prediction errors for human ages gradually increased indicated that advanced brain connection or activities emerges during development to boost evolutionary differences. Contrarily, the decreased prediction errors for macaque ages using human model indicated that mature macaque brain may ape young human at some extent. Overall, these findings suggest that the intrinsic evolutionary principles of the brain are effectively captured from a perspective along the timeline with the predictive model we developed.

Although many previous studies have used cortical expansion index to quantitatively characterize brain evolution (Hill et al., 2010), the findings about cortex expansion are inconsistent and even contradictory (Amlien et al., 2016; Chaplin et al., 2013; Foster et al., 2022). Thus, there is still lack of a stable index to quantify evolution differences in an intuitive perspective. Based on the cross-species prediction model, we proposed the concept of the brain cross-species age gap (BCAP) index to quantify brain evolution along with temporal axis. We observed that the BCAP is negatively correlated with visual sensitivity test while it is positively correlated with picture vocabulary test indicating BCAP indeed reflect the evolutionary behavioral phenotypes. To sum up, the BCAP may provide an object index to quantify cross-species brain development and evolution along with chronological axis.

### The human- and macaque-specific structural features for intra- and inter-species age prediction

An interesting finding for the prediction features is that the highest proportion of human- or macaque-specific features are primarily located in white matter tracts and gray matter brain areas, respectively. We also observed that disproportionate anatomical development exhibited in human and macaque brain during the adolescence. The human-specific features are closely related to brain development and mainly include white matter tracts of left arcuate fasciculus (AF), inferior fronto-occipital Fasciculus (IFOF) and superior longitudinal fasciculus I (SLF I) which are involved in language and complex motor tasks (Jacquemont et al., 2023; Rilling et al., 2011). The AF is an important language pathway connecting the key speech production region (Broca’s area) in the frontal lobe with the speech comprehension region (Wernicke’s area) in the posterior temporal lobe (Catani and Mesulam, 2008; Dick and Tremblay, 2012; Rilling et al., 2008). The lateralization of AF is reported to be closely associated with language evolution (Becker et al., 2022; Eichert et al., 2019). The macaque-specific features are mainly distributed in gray matter areas of the prefrontal cortex and motor cortex which are mainly involved saccade or visually-guided motor and general motor. In addition, some human- or macaque-specific features also found in gray matter areas and white matter tracts, respectively. The human-specific gray matter features are mainly localized in insula, basal ganglia and sensory cortex which play an important role in emotion, motor, and sensory information control (Chang et al., 2013; Draganski et al., 2008; Menon and Uddin, 2010). The macaque-specific white matter tracts features mainly include left superior longitudinal fasciculus II (SLF II) and right uncinate fasciculus which is related to working memory and memory retrieval (Janelle et al., 2022; Makris et al., 2005; Papagno et al., 2011). The analysis of human- and macaque-specific features revealed notable differences in the degree of development of regions associated with language and working memory functions during adolescence between humans and macaques.

### Evolution related brain areas and white matter tracts

The brain cross-species age gap was adopted to reveal evolution related brain areas and white matter tracts and identified similar findings with the human-specific features indicating disproportionate anatomical brain evolution along the chronological axis. The highly positively correlated white matter tracts are arcuate fasciculus (AF), optic radiation (OR) and right superior longitudinal fasciculus II (SLF II). Language is considered to be the predominant difference between human and other species during evolution. AF is involved in syntax and lexical-semantics processing and is reported to be closely associated with language evolution (Becker et al., 2022; Rilling et al., 2011). The AF is left lateralization and dorsal predominant in human while symmetrical and ventral predominant in macaques (Balezeau et al., 2020a; Eichert et al., 2019; Rilling et al., 2008). In addition, evolution related right SLF II connecting parietal cortex and frontal cortex was found. A previous study using task fMRI identified an attention-specific brain area within temporoparietal junction in human (Patel et al., 2015). Thus, the evolution associated right SLF II found in this study may be the underlying white matter pathway for attention.

Moreover, evolution related gray matter brain areas including ventrolateral, medial, centrolateral, dorsomedial, and dorsolateral prefrontal cortex were also uncovered. These frontal cortical areas are involved in high-order cognitive functions including cognitive control and executive, decision making, plan, reasoning and so on (Hiser and Koenigs, 2018; Ray and Zald, 2012; Salzman and Fusi, 2010). The prefrontal cortex has been widely reported during evolution with the largest expansion, late myelination and structural asymmetry in human compared to other nonhuman primates (Li et al., 2017; Miller et al., 2012; Smaers et al., 2013). In addition to evolution associated gray matter areas and white matter tracts, we also found some gray matter areas of putamen, amygdala and subgenual anterior cingulate cortex and white matter tracts of Forceps Minor (FM) and OR are negatively correlated with the brain cross-species age gap. These gray matter areas and white matter tracts mainly participate in motor, visual and emotion (Gray et al., 2002; Murray, 2007). The negative correlation may suggest that these brain areas or white matter tracts are more predominant in macaque than that in human. Taken together, using the brain cross-species age gap index, we identify the evolution associated brain areas and white matter tracts widely reported in previous studies and discovered that the human brain shows a greater proportion of development in language and higher-order cognitive functions through cross-species comparisons of brain development along with chronological axis, which demonstrated that the brain cross-species age gap is an effective index for future studies to quantify brain evolution between species.

In conclusion, we proposed to embed the brain anatomy of human and macaque in the chronological axis for enabling the cross-species comparison using brain structure based cross-species age prediction model on brain development. We demonstrated that prediction model has good performances for cross-species age prediction and the trained macaque model to predict human ages outperform the trained human model to predict macaque ages. Moreover, we proposed the index of the brain cross-species age gap (BCAP) to quantify brain evolution along with temporal axis and identified evolution associated gray matter areas and white matter tracts consistent with previous reports. Overall, we introduced a novel protocol and index to investigate and quantitatively characterize brain evolution and discovered disproportionate anatomical development and evolution involving language pathways and cognitive control for human and working memory function for macaque, which may promote the development of comparative neuroscience or neuroimaging.

## Materials and methods

### Human subjects and MRI data acquisition

A total of 370 healthy human subjects (170 males/200 females, age range of 8-14 years, mean and standard deviation of 11.1 ± 1.8 years) with high quality structure magnetic resonance imaging (sMRI) and diffusion MRI (dMRI) were included in this study. All the MRI data were scanned using Siemens 3T Tim Trio MRI scanner with 32-channel head coil and simultaneous multi-slice echo planar imaging sequence. Sagittal 3D T1-weighted images were acquired using the following parameters: time repetition (TR) = 2500 ms, time echo (TE) = 2.22 ms, flip angle = 8°, voxel resolution: 0.8×0.8×0.8 mm^3^. The acquisition parameters for dMRI were: 185 directions on 2shell of b = 1500 and 3000 s/mm^2^, along with 28 b = 0 s/mm^2^, TR/TE = 3230/89.2 ms, flip angle = 78°, voxel resolution: 1.5×1.5×1.5 mm^3^, 92 slices. The detailed MRI scanning parameters of the MRI data can be found in a previous study (Somerville et al., 2018). In addition, the behavioral phenotypes including Behavioral Inhibition System and Behavioral Activation System Scale, Child Behavior Checklist, Visual Sensitivity Test, and Picture Vocabulary Test were also assessed for each individual. The used public MRI and behavioral phenotypes data were accessed through the Human Connectome Project-Development (HCP-D: http://www.humanconnectome.org). In order to approximately match the ages of the used macaque data in this study, only 370 subjects with age from 8-14 years from HCP-D were finally analyzed. All subjects were given informed written consents approved by the Institutional Review Board.

### Macaques and MRI data acquisition

The structural and diffusion MRI data of 181 macaque monkeys (Macaca mulatta, 89 males/92 females, age range from 2-4 years, mean and standard deviation = 2.8 ± 0.6 years) were used in this study. All the macaque MRI datasets were acquired at University of Wisconsin-Madison (UW-Madison), of which 42 macaques were scanned using GE DISCOVERY_MR750 3.0T MRI, and the other 139 macaques were scanned using GE Signa EXCITE 3.0T MRI. All the experimental procedures were approved by the University of Wisconsin Institutional Animal Care and Use Committee in compliance with the Guide for the Care and Use of Laboratory Animals published by the US National Institutes of Health. Before scanning, all the animals were anesthetized with the following procedures: ketamine, medetomidine, ketamine maintenance anesthesia. Macaques were scanned in the sphinx position with the nose pointing into the scanner after approximately 30 minutes from first ketamine administration. The physiological features including heart rate and oxygen saturation were monitored and recorded at minimum every 15 minutes during scanning. Heated water bags, bottles, or pads and towels, blankets, and bubble wrap were used to maintain body temperature during imaging. The scanning parameters of dMRI with GE DISCOVERY_MR750 3.0T were follows: 12 directions with b ≈ 1000 s/mm^2^, TE/TR = 94.3/6100 ms, flip angle = 90°, voxel resolution = 0.5469 × 2.5 × 0.5469 mm^3^. The structural T1 images scanning parameters with GE DISCOVERY_MR750 3.0T were: TR/TE = 11.4/5.412 ms, flip angle = 10°, voxel resolution: 0.2734 × 0.5 × 0.2734 mm^3^. For GE Signa EXCITE 3.0T MRI scanner, the dMRI scanning parameters were: 12 directions with b ≈ 1000 s/mm^2^, TE/TR = 77.2/10000 ms, flip angle = 90°, voxel resolution = 0.5469 × 2.5 × 0.5469 mm^3^. The structural T1 images scanning parameters with GE Signa EXCITE 3.0T MRI were: TR/TE = 8.648/1.888 ms, flip angle = 10°, voxel resolution: 0.2734 × 0.5 × 0.2734 mm^3^. The detailed MRI scanning parameters are available online (http://fcon_1000.projects.nitrc.org/indi/PRIME/uwmadison.html) and the detailed scanning parameters can refer to a previous study (Young et al., 2017). All the information for human and macaque was presented in Table 1.

**Table 1.** Demographic information for human and macaque.

| Subjects | Human<br>(8-14 years) | Macaque<br>(2-4 years) |
| --- | --- | --- |
| Number of subjects | 370 | 181 |
| Gender (male: female) | 170 / 200 | 89 / 92 |
| Age (mean ± SD) | 11.07 ± 1.75 | 2.75 ± 0.57 |
| Male age (mean ± SD) | 10.89 ± 1.61 | 2.62 ± 0.51 |
| Female age (mean ± SD) | 11.23 ± 1.85 | 2.87 ± 0.60 |

### Human and macaques structural MRI data preprocessing

The gray matter volume (GMV) of human and macaque was calculated using voxel-based morphology (VBM) with SPM8 package (http://www.fil.ion.ucl.ac.uk/spm). The VBM analyses for human subjects using a fully automatic procedure included the following steps: the T1 image was first segmented into gray matter, white matter, and cerebrospinal fluid; the segmented images were then transformed to MNI space using high dimensional DARTEL normalization and were modulated to account for volume changes. For macaque VBM analysis, the main procedures as follows: the skull of the structural T1 image was first manually stripped and then was rotated to match the orientation of the INIA19 template (Rohlfing et al., 2012); the skull-stripped T1 image was then automatically segmented into gray matter, white matter and cerebrospinal fluid and was registered to the INIA19 template using DARTEL-normalization; mean template images of gray matter, white matter and cerebrospinal fluid were created and each individual gray matter volume was transformed into the new mean gray matter template image. The details for calculating the human and macaque GMV have been described in our previous studies (Wang et al., 2018; Wang et al., 2017).

### Human and macaques dMRI data preprocessing

The dMRI data for both human and macaque were preprocessed using the FSL software (http://www.fmrib.ox.ac.uk/fsl). The linear portion of eddy currents and head motions of the diffusion MRI were corrected by registering all images to the b0 image using affine transformation to minimize gradient coil eddy current distortions using *eddy_correct* function. Next, the indices of FA, MD, AD, and RD were calculated to quantify the white matter microstructural integrity and/or myelination. Finally, all the FA, MD, AD, RD maps were registered to template images (MNI152 template for human and INIA19 template for macaque) for further analyses.

### Harmonization for GMV and DTI-derived indices

Given that the differences in imaging protocols, scanning parameters and scanner manufacturers may affect the reliability of MRI-derived measurements (Alexander et al., 2001; Correia et al., 2009; Giannelli et al., 2009; Zhan et al., 2010), we conducted harmonization processing on macaque datasets to ensure the consistency for the following analyses. The harmonization of macaque datasets was performed using ComBat method which has been widely applied in previous studied to mitigate bias induced by scanning differences (Eshaghzadeh Torbati et al., 2021; Fortin et al., 2017; Johnson et al., 2007).

### Brain structure based cross-species age prediction model

In this study, we developed a brain structure based cross-species age prediction model to quantitatively characterize evolution patterns. The indices of GMV, FA, MD, AD and RD were used as features to construct prediction models for humans and macaques. For cross-species prediction, we used homogenous brain gray matter and white matter atlases to extract same features for human and macaque. The brain gray matter and white matter were parcellated into 92 sub-regions or 42 sub-tracts in both human and macaque using regional map (RM) atlas and XTRACT cross-species atlas, respectively (Bezgin et al., 2017; Warrington et al., 2020). For each human subject or macaque, a total of 260 features (168 features for white matter tracts (FA, MD, AD, RD for 42 tracts) and 92 features for GMV) were obtained. To select the ultimate features for prediction, two steps were adopted to determine the optimal features for the final intra- and inter-/cross-species prediction. First, the Pearson’s correlation analyses between all the 260 features and ages across all human subjects or all macaques were performed and the features showing significant correlations with *p* < 0.01 were used for prediction. Second, we separately trained the prediction models in humans and macaques using the selected features as stated above and the validity of prediction models were tested using 10-fold cross-validation. We used nine tenths features for training and the remaining one tenth features for testing and this procedure was repeated 100 times. Since the selected features in the 100 times were different, we determined the final features for intra- and inter-/cross-species prediction using two criteria. The first is to extract the features that are present in all the 100 repetitions. Through this way, a total of 62 features for macaque model and a total of 225 features for human model were selected. The second way is to extract the features showing minimum mean absolute error (MAE) for prediction in one of the 100 times repetitions and finally a total of 117 features for macaque and 239 features for human were acquired. The selected features through the above two methods were both used for the following intra- and inter-/cross-species prediction to exclude the effects of different number of features. In addition, to further exclude the influences of unequal number of features in human and macaque, we only selected the top 62 features of human and macaque to test the prediction model performances.

With the selected features, four prediction models were constructed: human brain structural features predict human ages; macaque brain structural features predict macaque ages; human brain structural features predict macaque ages and macaque brain structural features predict human ages. Three sets of features (62 macaque features and 225 human features; 117 macaque features and 239 human features; 62 macaque features and 62 human features) were used to test the prediction models for cross-validation and to exclude effects of different number of features in human and macaque. For all these types of prediction models, 10-fold cross-validation with nine tenths features for training and the remaining one tenth features for testing were performed again to obtain the final prediction results. The prediction performances were evaluated by calculating the Pearson’s correlation and MAE between actual ages and predicted ages.

To quantify potential evolutionary differences for the inter-/cross-species prediction models, Pearson’s correlation coefficients between the actual ages and the brain age gap (|Δ _brain age_|) defined as absolute value of actual age minus predict age were calculated for both human and macaque.

To determine the features distributions of the prediction models in human and macaque, we classify the used features of different measures (FA, MD, AD, RD, GMV) into shared/common, human-specific and macaque-specific features. For each measure, the percentage of the three different types of features was computed. In addition, for each type of the features, the top 5 brain areas or white matter tracts and the corresponding correlation coefficients with ages were shown.

In addition, to test whether sex affects prediction results, we divided both human and macaque participants into male and female groups. The prediction models were separately trained in male and female and were applied to predict the ages of the corresponding groups’ subjects. For inter-/cross-species prediction, the trained human or macaque prediction models in male or female were applied to predict the macaque or human ages in male or female, respectively.

### The brain cross-species age gap associated brain areas, white matter tracts and behavioral phenotypes

To better quantitatively characterize brain evolution, we proposed the concept of the brain cross-species age gap (BCAP) defined as percentile rank differences which has been demonstrated to be easier to interpret a score or a data point within a data set (Crawford et al., 2009). To calculate BCAP, each human subject’s predicted age using human prediction model (denote as P-age _human-human_) was first obtained. Then, the same human subject’s predicted age using monkey prediction model (denote as P-age _monkey-human_) was also calculated. Next, the percentile rank of P-age _human-human_ and P-age _monkey-human_ were computed across all the subjects’ predicted P-age _human-human_ and P-age _monkey-human_, respectively. Finally, the BCAP was defined as the percentile of P-age _human-human_ minus the percentile of the P-age _monkey-human_.

To test whether BCAP could reflect behavioral phenotypes, Pearson’s correlation analyses between BCAP and 40 behavioral phenotypes including Early Adolescent Temperament Questionnaire, Edinburgh Handedness Questionnaire, Autism Spectrum Rating Scales, Behavioral Inhibition System and Behavioral Activation System Scale, etc. across all the human subjects were performed with actual ages as covariates. The significant level was determined using false family discovery (FDR) method with P < 0.05.

Finally, with the obtained BCAP for each human subject, Pearson’s correlation analyses between BCAP and GMV of each brain area, FA, MD, AD, RD values of each white matter tract across all the human subjects were performed with actual ages as covariates to identify evolution associated brain areas and white matter tracts. The significantly associated brain areas and white matter tracts were determined using FDR method with P < 0.05. The top 3 white matter tracts in FA and the top 5 gray matter brain areas showing the highest and lowest correlation values with ages were shown.

## Acknowledgement

This study was supported by the National Natural Science Foundation (62176044), Natural Science Foundation of Yunnan Province (major basic research project: grant number: 202102AA100053) and Yunnan Fundamental Research Projects (202201BE070001-004).

## Author contributions

J.W and C.C contributed to the conception and design of the study. Y.L, Q.S and S.Z contributed to the downloading and analysis of data; Y.L, J.W and C.C wrote and edited the manuscript. All the authors discussed the paper.

## Compliance with Ethical Standards

This study used a public dataset which obtained the ethical improvement.

## Supplementary information

**Table supplement 1.** The information of 62 selected features of macaque. The first column shows the four brain parameters, the second column shows selected white matter tracts and gray matter regions within the brain parameters, and the third column shows the R-value using Pearson’s correlation between the second column and ages.

**Table supplement 2.** The information of 225 selected features of human. The first column shows the four brain parameters, the second column shows selected white matter tracts and gray matter regions within the brain parameters, and the third column shows the R-value using Pearson’s correlation between the second column and ages.

**Table supplement 3.** The information of 117 selected features of macaque. The first column shows the four brain parameters, the second column shows selected white matter tracts and gray matter regions within the brain parameters, and the third column shows the R-value using Pearson’s correlation between the second column and ages.

**Table supplement 4.** The information of 239 selected features of human. The first column shows the four brain parameters, the second column shows selected white matter tracts and gray matter regions within the brain parameters, and the third column shows the R-value using Pearson’s correlation between the second column and ages.

**Figure supplement 1.**
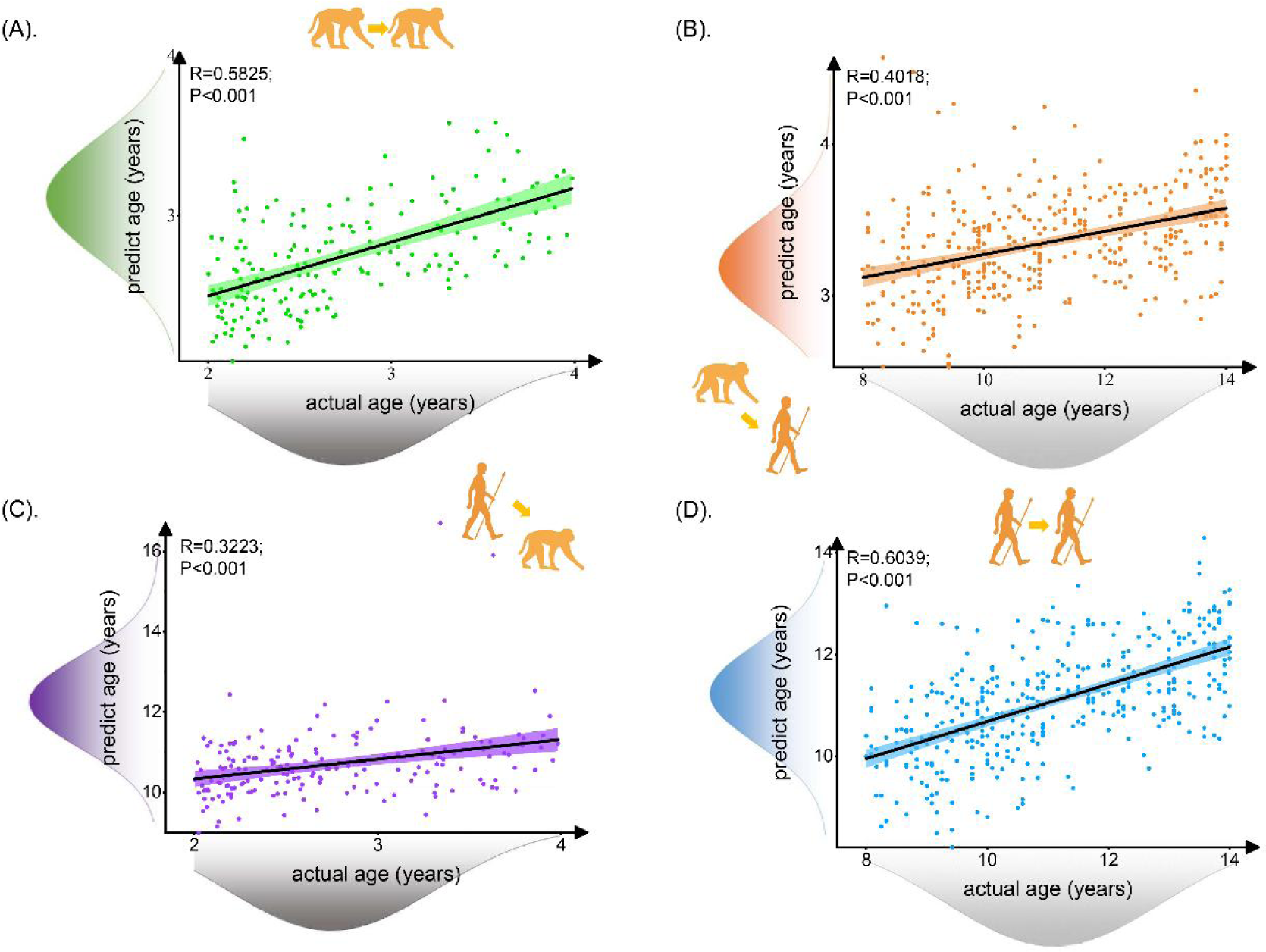
The prediction results of intra- and inter-species using prediction model with 117 selected macaque features and 239 selected human features. Each dot depicts data from an individual participant. The width of the curve denotes the 95% confidence interval around the linear fitting curve (black line). Normal distribution of predicted ages and actual ages expressed on the left and bottom of scatterplot. The prediction model could well predict intra- and inter-species ages.

**Figure supplement 2.**
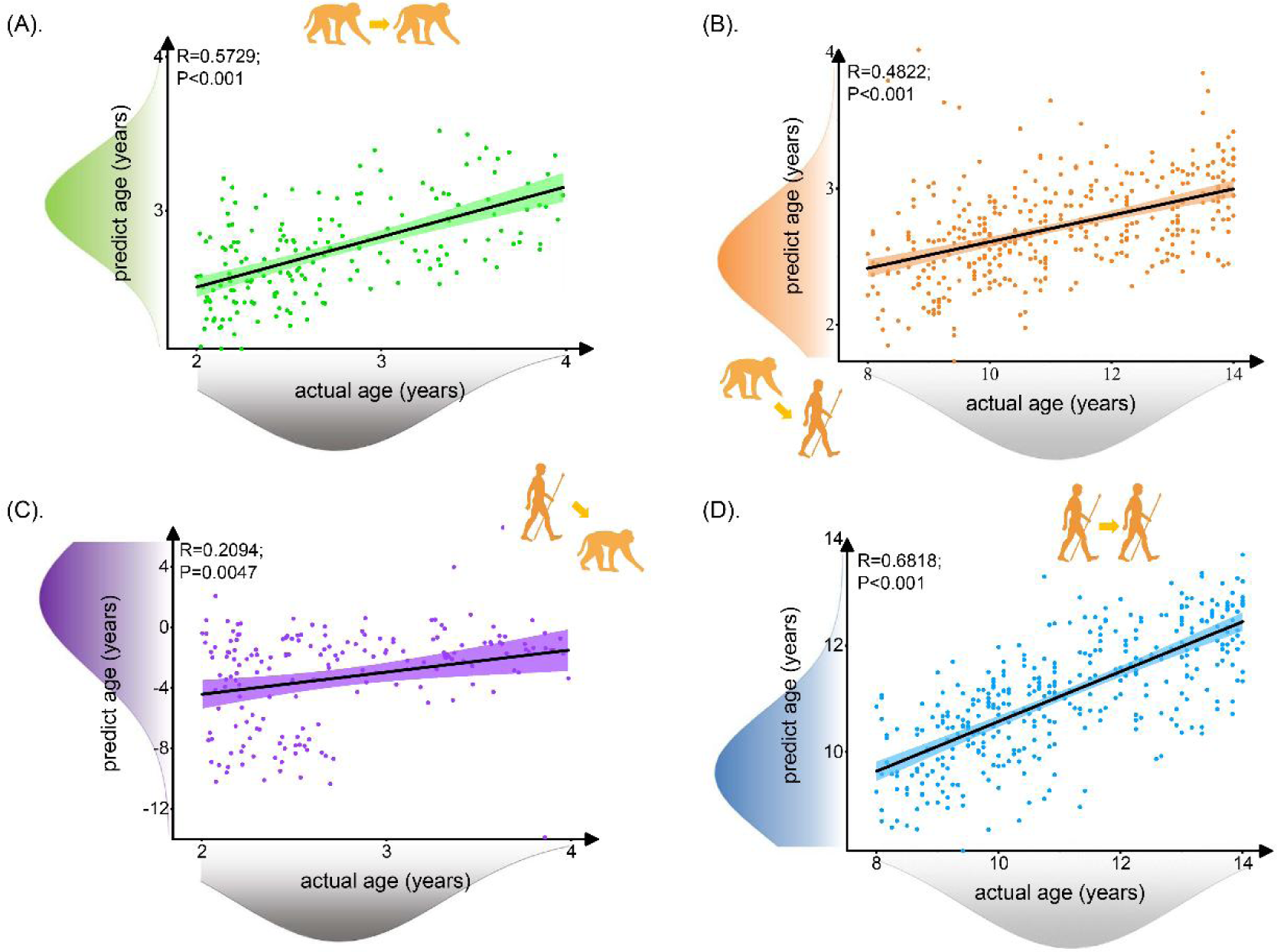
The prediction results of intra- and inter-species using prediction model with 62 selected macaque features and 62 selected human features. Each dot depicts data from an individual participant. The width of the curve denotes the 95% confidence interval. Normal distribution of predicted ages and actual ages expressed on the left and bottom of scatterplot. The prediction model could well predict intra- and inter-species ages.

**Figure supplement 3.**
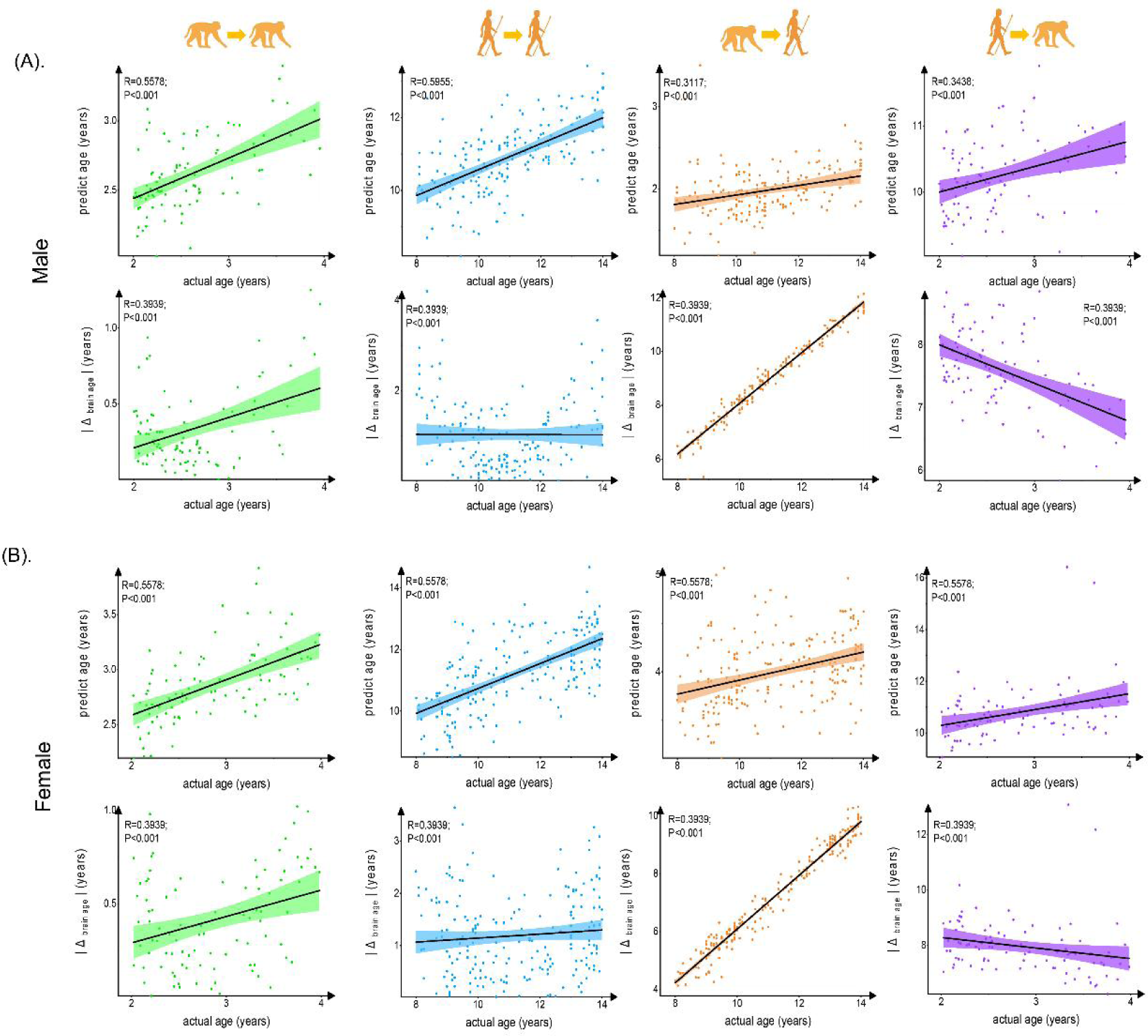
The prediction results of intra- and inter-species in male and female. Each dot depicts data from an individual participant. The width of the curve denotes the 95% confidence interval. The prediction in intra- and inter-species observed significant and no evolution differences between male and female. (A). The prediction results in male. (B). The prediction results in female.

**Figure supplement 4.**
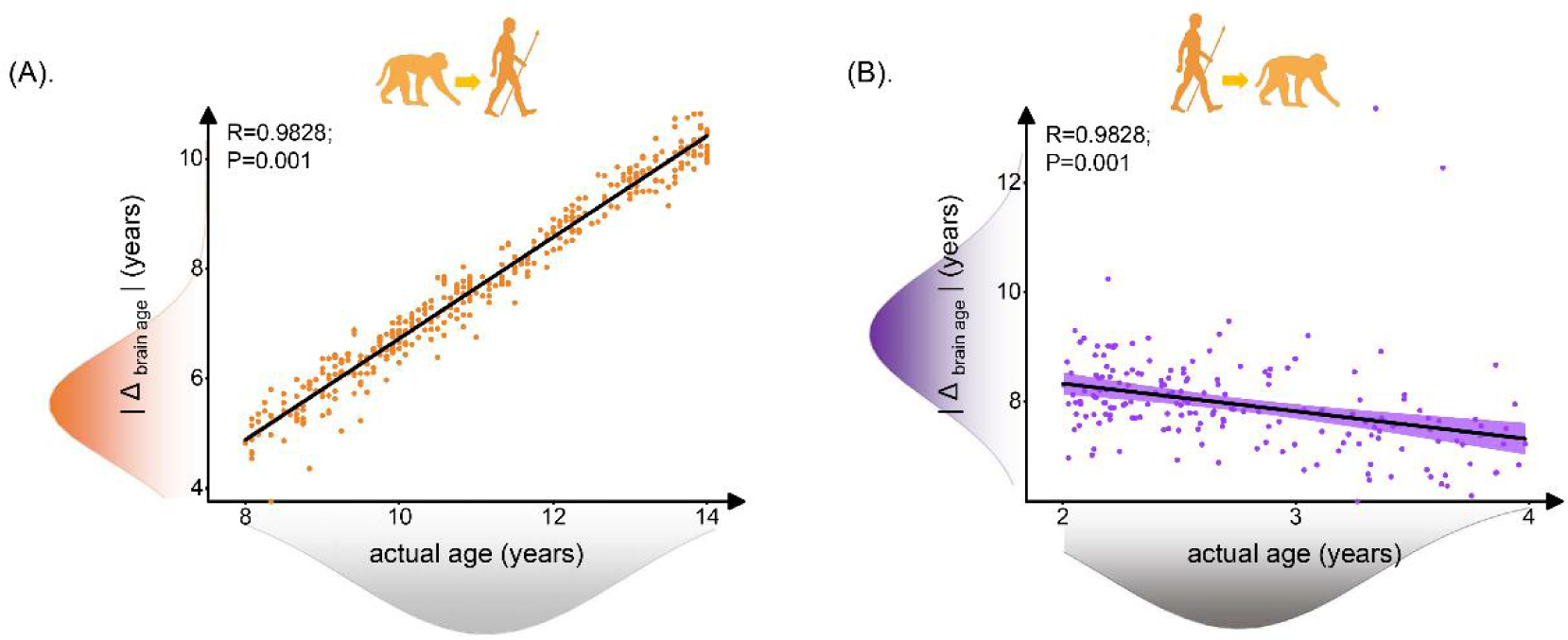
Relationship between brain age gap (|Δ _brain age_|) and actual ages in human and macaque with 117 selected macaque features and 239 selected human features. Each dot depicts data from an individual participant. The width of the curve denotes the 95% confidence interval. (A). Positive relationship between | Δ _brain age_ | and actual age in macaque model predicted human ages (Pearson’s correlation: R = 0.9828, P = 0.001, MAE = 3.3545). (B). Negative relationship between | Δ _brain age_ | and actual age in macaque model predicted human ages (Pearson’s correlation: R = −0.3318, P < 0.001, MAE = 5.2055).

**Figure supplement 5.**
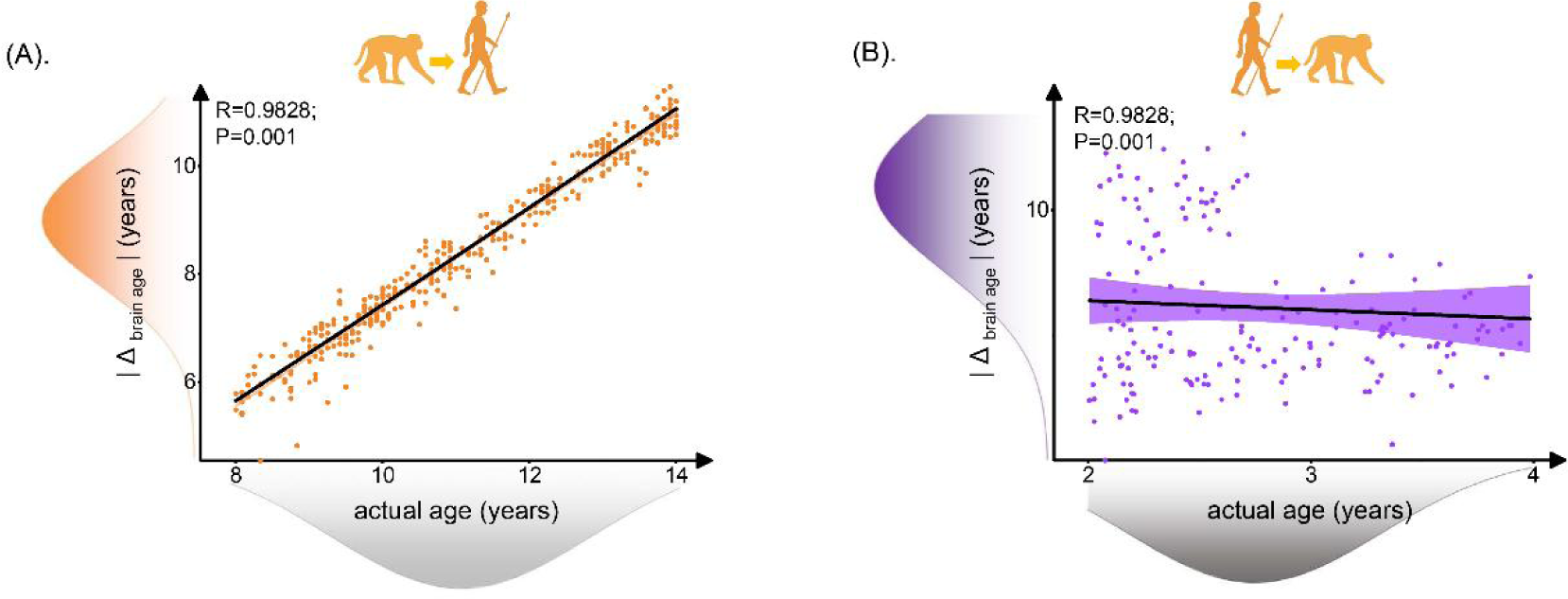
Relationship between brain age gap (|Δ _brain age_|) and actual ages in human and macaque with 62 selected macaque features and 62 selected human features. Each dot depicts data from an individual participant. The width of the curve denotes the 95% confidence interval. (A). Positive relationship between | Δ _brain age_ | and actual age in macaque model predicted human ages (Pearson’s correlation: R = 0.9814, P < 0.001, MAE = 2.7123). (B). Negative relationship between | Δ _brain age_ | and actual age in macaque model predicted human ages (Pearson’s correlation: R = −0.054, P = 0.4704, MAE = 3.4821).

**Figure supplement 6.**
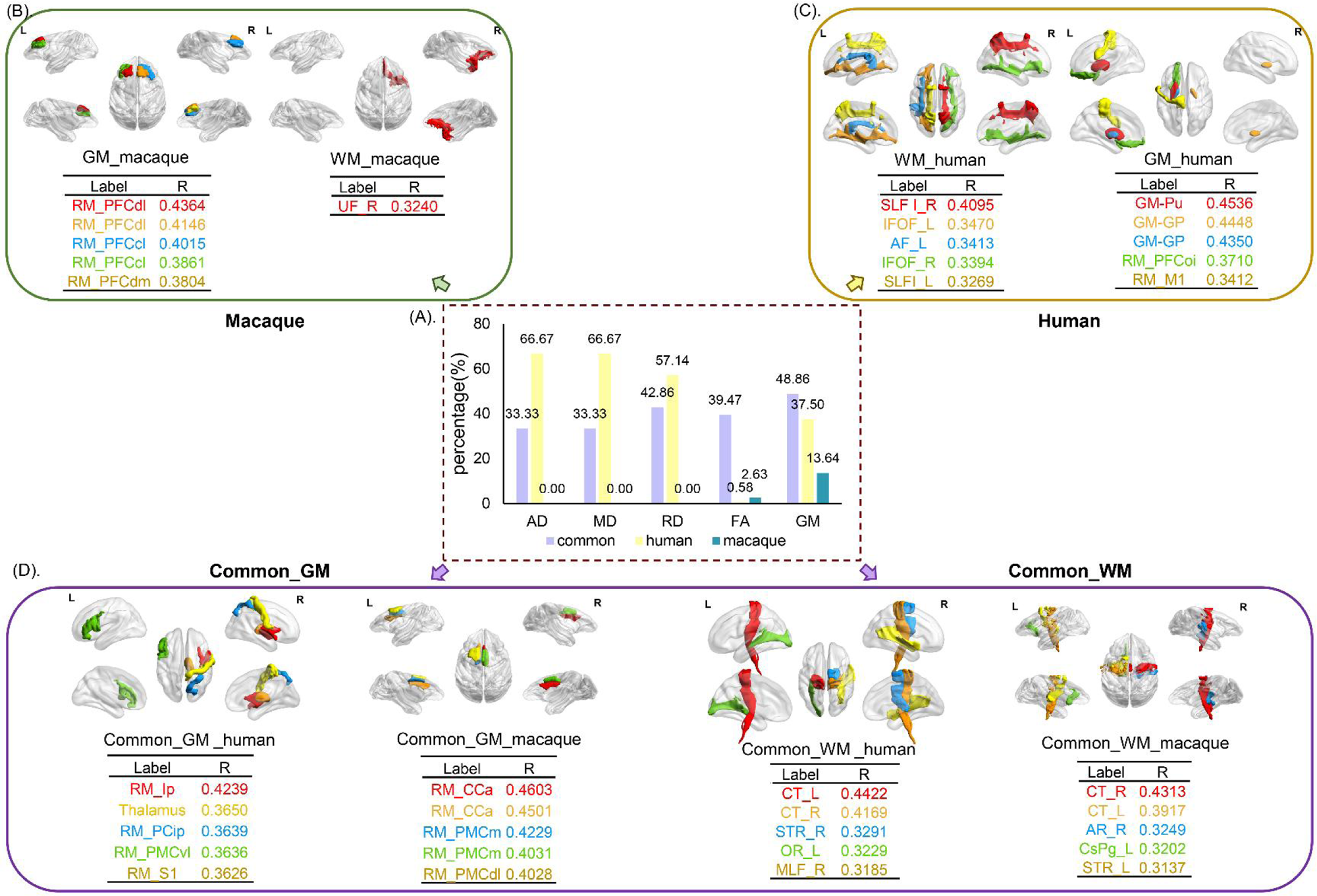
The distribution of features with 117 selected macaque features and 239 selected human features. The selected features were analyzed based on five parameters (GMV, FA, MD, RD and AD) and three groups (human-specific, macaque-specific and common features in human and macaque). (A). The percentage of each group in each parameter. The macaque-specific features are only located in FA and GMV. (B),(C),(D). The top five features of macaque-specific, human-specific and common in human and macaque in gray matter and white matter tracts except that only one macaque-specific features in white matter tracts. FA: fractional anisotropy; MD: mean diffusivity; RD: radial diffusivity; AD: axial diffusivity; GMV: gray matter volumes.

**Figure supplement 7.**
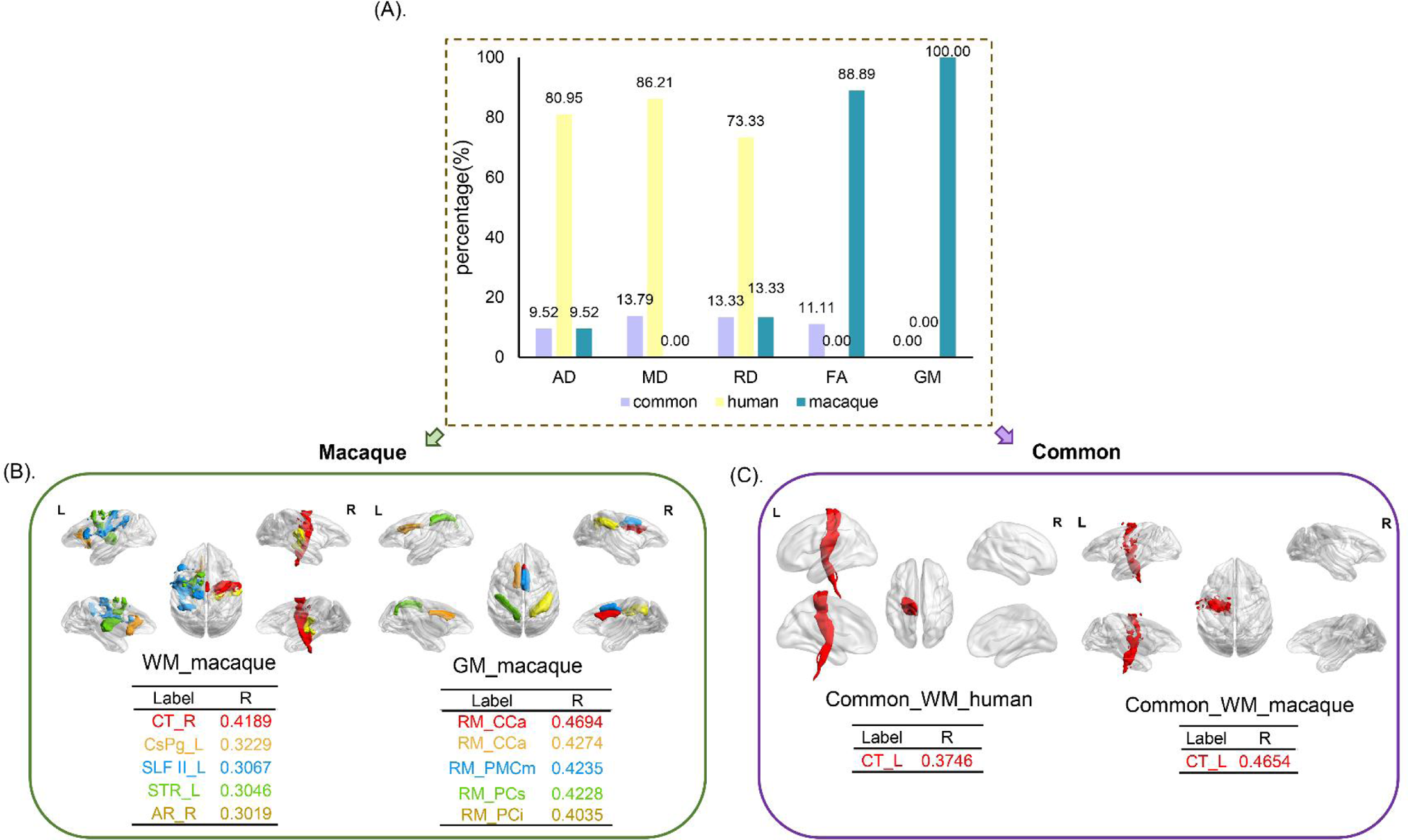
The distribution of features with 62 selected macaque features and 62 selected human features. The selected features were analyzed based on five parameters (GMV, FA, MD, RD and AD) and three groups (human-specific, macaque-specific and common features in human and macaque). (A). The percentage of each group in each parameter. The macaque-specific features are only located in FA and GMV. (B),(C). The top five features of macaque-specific in gray matter and white matter tracts and only one common feature in human and macaque. FA: fractional anisotropy; MD: mean diffusivity; RD: radial diffusivity; AD: axial diffusivity; GMV: gray matter volumes.

**Figure supplement 8.**
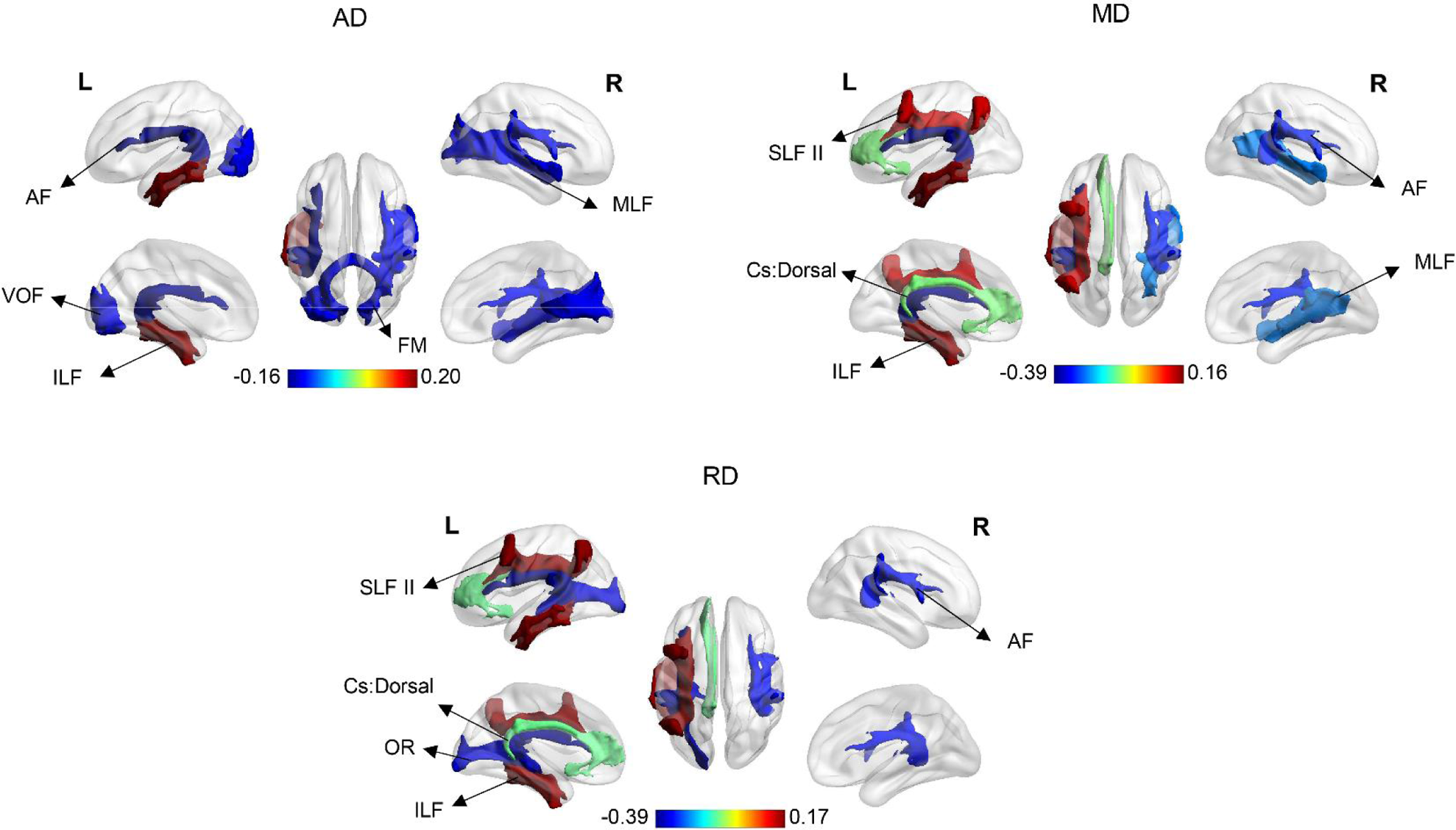
The top three and last three features associated with E-age gap in MD, RD and AD. AF: arcuate fasciculus; VOF: vertical occipital fasciculus; ILF: inferior longitudinal fasciculus; FM: forceps major; MLF: middle longitudinal fasciculus; SLF II: superior longitudinal fasciculus II; Cs:Dorsal: cingulum subsection: Dorsal; OR: optic radiation.

